# Adaptation to nighttime light via gene expression regulation in *Drosophila suzukii*

**DOI:** 10.1101/2025.02.17.638639

**Authors:** Natsumi Takenaka, Yuma Takahashi

## Abstract

Urbanization causes environmental changes like habitat loss, fragmentation, and pollution, which reduce biodiversity. Urban organisms face stressors, such as heat islands, air and water pollution, and anthropogenic noise, all of which can disrupt their development, behavior, and physiology. While some species adapt to urban environments, their responses and the role of evolution in urbanization are limited, as most studies focus on phenotypic traits. Artificial light at night (ALAN), a common urban stressor, disrupts behaviors and physiological processes, including circadian rhythms, sleep, and reproduction. The present study examined ALAN’s effects on body size, survival, activity rhythms, and gene expression in *Drosophila suzukii*, a species found in urban and rural habitats. ALAN reduced wing and thorax sizes regardless of sex and origin, decreased survival in rural populations, and increased it in urban populations. ALAN elevated overall activity, especially in the early night, while urban females displayed reduced sensitivity regarding activity and sleep. The circadian rhythm length was disrupted in rural populations but not in urban populations. Transcriptomic analysis revealed ALAN-induced gene expression changes, particularly in urban females, with photoreceptor- and circadian rhythm-related genes responding differently between urban and rural populations. These results indicate that urban populations have evolved adaptive mechanisms to counter ALAN’s effects, likely mediated through gene regulation. This study highlights ALAN’s impact on diverse traits and its potential for adaptive evolution in urban environments. Evolutionary adaptations in traits related to urban stress responses may enhance the ecological success of *D. suzukii* in urban habitats.

## Introduction

Urban populations are expected to grow substantially in the coming decades, accompanied by a rapid expansion of urban areas (Huang et al. 2019). This urban expansion is anticipated to have diverse ecological impacts, such as habitat fragmentation, isolation, and loss (Liu et al. 2016). These changes often lead to the reduction or extinction of species and populations, contributing to the biodiversity decline (Xu et al. 2018; Theodorou et al. 2021). Urbanization was also suggested to be linked to the ongoing issue of insect population declines, i.e., insect apocalypse (Vaz et al. 2023).

In addition to fragmentation, urban habitats undergo abiotic changes. Organisms in urban areas are exposed to multiple stresses caused by increased artificial light at night (ALAN, light pollution), heat islands, water and air pollution, and anthropogenic noise (noise pollution). These stresses are thought to affect the organisms’ development, behavior, and physiology and are potential factors that may alter the demographic dynamics of species and populations (Schmidt et al. 2020). While some species disappear due to rapid urban environmental changes, others thrive in urban environments (Hahs et al. 2023). Therefore, tolerance to urban stress is an essential factor for the survival of individuals in urban environments. For example, plasticity in heat stress tolerance has been suggested to contribute to expanding the distribution range to urban areas (Sato and Takahashi 2022). However, plasticity does not always lead to an adaptive response to the new environment; it can also result in nonadaptive or maladaptive responses, resulting in harmful effects. These facts indicate that the plastic or developmental response to urban stress could inhibit adaptation and survival in urban environments (Tüzün et al. 2017; Campbell-Staton et al. 2021). Therefore, it is essential to carefully examine the phenotypic effects of urban stress on survival and reproduction to accurately understand and predict the impact of urbanization on organisms.

Recently, evidence for contemporary adaptive evolution to the urban environment has been increasingly documented across various taxa (Johnson and Munshi-South 2017; Rivkin et al. 2019). The evolution of tolerance to urban stress is suggested to enable species to inhabit cities. However, many studies focusing on adaptive evolution in cities failed to distinguish between plastic phenotypic responses to environmental factors and evolutionary changes in phenotypes; most previous studies measured phenotypes based on individuals collected from the field. Common garden or transplant experiments are required to distinguish genetic from nongenetic changes. The simultaneous quantification of genetic and nongenetic changes in each trait is essential to understanding the overall response to ongoing urbanization and predicting the ecological responses of organisms in urban areas.

Previous studies have explored how urban stress affects phenotypic traits crucial for fitness and survival (Senar et al. 2014; Multini et al. 2019). However, this trait-based approach limits the ability to fully identify traits influenced by urban stress or shaped by contemporary urban development. In contrast, genome and transcriptome analyses can comprehensively detect the potential effects of urbanization (Salmón et al. 2021; Fukano et al. 2023; Minias 2023). For instance, transcriptome analysis can identify genes with altered expression patterns due to urbanization stress and highlight evolutionary differences between sites at varying urbanization levels. Combining omics data with phenotypic traits is essential for understanding biological responses to urban stress.

ALAN is an environmental change associated with rapid urbanization (Cinzano et al. 2001; Kyba et al. 2015; Falchi et al. 2016). Recent studies have revealed that it significantly impacts the diurnal activity of organisms across various taxa, including insects, fish, birds, and mammals. ALAN disturbs circadian rhythms and alters a range of traits, reducing immune function (Durrant et al. 2015), affecting development (Durrant et al. 2018a), disrupting endocrine functions (Russart and Nelson 2018), and reducing sleep (Kolbe et al. 2021) and daytime activities (Sato and Takahashi 2022). Thus, ALAN is thought to significantly impact the survival and reproduction of organisms in urban areas. However, while many studies are available on birds and mammals (Hoffmann et al. 2018; Jiang et al. 2020), distinguishing whether the response to ALAN represents a plastic or evolutionary response is often challenging.

Here, we examined the direct effects of ALAN on the morphological and life-history traits and gene expression patterns in *Drosophila suzukii*, an agricultural pest species. We also compared these effects between urban and rural populations to identify their evolutionary differences. As with other drosophila species, this species can be maintained in the laboratory, and its plastic and evolutionary responses to urbanization stress can be quantified separately using common garden experiments.

## Materials and Methods

### Study species

The spotted wing drosophila *D. suzukii* is a close relative of *D. melanogaster*. It originated in Asia and has recently invaded most of Europe and South America (Lee et al. 2011). *D. suzukii* lays eggs in thin-skinned soft fruits, such as strawberries, blackberries, raspberries, and cherries. It is widely regarded as one of the most important agricultural pest species. The flies can inhabit urban and rural areas. In Japan, they primarily feed on cherry fruits and bayberries, and adults mainly emerge in spring and autumn.

### Sampling and strains

From 2020 to 2022, we collected ripe cherry fruits as potential host fruits from urban and rural areas in the Kanto region of Japan (Table S1). The sampling points were at least 5 km apart. Isofemale *D. suzukii* lines were established from a single female and male that emerged from fruits collected at the same locations. The flies were reared for at least a year under constant conditions (12L12D, 22°C) to eliminate genetic variation within lines and the environmental and maternal effects. They were maintained in vials with food medium Formula 4–24 Instant Drosophila Medium (Carolina Biological Supply Company) and transferred to fresh vials approximately every 3 weeks.

### Light simulation and obtaining test individuals

Ten males and 20 females were put into fresh vials with food medium and allowed to mate and oviposit for 48 h. Eggs laid in a vial were maintained under a constant environment at 22°C in an incubator. In the control group, the vials were exposed to a 12 h light/dark cycle (control, 2,500 Lx in daytime and 0 Lx in nighttime), while in the treatment condition, they were exposed to a 12 h light/dim light cycle (ALAN treatment, 2,500 Lx in daytime and 10 Lx in nighttime, with light-on at 06:00 and light-off at 18:00). Bright and dim lights were typical white LED light. Mated flies younger than 5 days old were collected and used in the following experiments.

### Body size

Body size was measured according to previous studies (Lack et al. 2016) using adults preserved in 0.9% saline solution containing 70% ethanol to prevent the samples from shrinking. Measurements were taken under a Leica M 165 FC stereomicroscope and analyzed using Leica Application Suite morphometric software (LAS X; Leica, Wetzlar, Germany). The thorax length was measured as the distance between the base of the anterior humeral bristle and the posterior tip of the scutellum. The wing length was measured as the straight-line length connecting the intersection of the anterior cross-vein and L4 longitudinal vein to the point where the L3 longitudinal vein intersects the wing margin.

### Survival rate

One- to two-day-old presumably mated individuals were sexed and transferred to fresh vials. Their density was standardized to 10 individuals per vial. They were maintained in the same light conditions as they developed until they became adults. Individuals in a vial were transferred to a fresh vial every 3–5 days. The number of surviving individuals was recorded at every transfer. Mixed-effects Cox proportional hazards models, including light treatment and urbanization level as fixed effects and vial and isofemale line as random effects, were used to analyze survival, using the *coxme* function of the “coxme” package in R.

### Activity level, sleep-wake behavior, and circadian rhythm detection

Adult activity was monitored using the Locomotor Activity Monitor (LAM, TriKinetics, Waltham, MA, USA). Five females or five males were transferred into a glass tube (φ25 mm). The tube was sealed at the food end with a tube cap, and the opposing end was sealed with a sponge plug to allow for air exchange. The tubes were mounted on the LAM, which characterized the movement pattern in each tube using infrared beams. All experimental procedures were performed at 22°C in an incubator. Under LD conditions, environmental stimuli are accepted through the functional eye, directly regulating the activity (Schlichting et al. 2015). Therefore, flies were monitored for 60 h at 1-min intervals under constant dark (DD) conditions to investigate endogenous rhythms. Data collected from 0 to 12 h after the experiment onset were removed because they were not considered accustomed to the experimental environment. Therefore, data collected from 6 h to less than 18 h after the experiment onset were analyzed. Principal component analysis (PCA) was performed using data on a log-transformed locomotion counter per hour (i.e., 24 variables) to extract the indicator of the daily activity pattern. Sleep was defined as uninterrupted behavioral inactivity of 5 min following a previous study (Karen Ho and Sehgal 2000). Data were collected using TriKinetics software. We analyzed the activity count using the Rethomics framework to explore the effect of ALAN on sleep behavior and the length of the circadian rhythm period (Geissmann et al. 2019). We used the χ^2^ periodogram method to predict the periodicity.

### RNA extraction

Ten males and 20 females were put into fresh vials filled with food medium and allowed to mate and oviposit for 48 h. Eggs laid in a vial were maintained under control and treatment conditions at 22°C in an incubator. Mated flies under 3 days old were collected and used for RNA extraction. Flies were transferred into a 0.8 ml microcentrifuge tube under CO_2_ anesthesia at an approximate zeitgeber time of 11 and immediately stored at −80°C. Their heads were removed with forceps, and 10 heads were simultaneously homogenized using a pestle. Total RNA was extracted using the Maxwell 16 LEV Plant RNA Kit with the Maxwell 16 Research Instrument (Promega) following the manufacturer’s protocol. Electrophoresis on a 1% agarose gel was performed to check for extraction quality. The RNA was eluted with nuclease-free water, and RNA concentrations were estimated using a Qubit 2.0 Fluorometer (Invitrogen). RNA purity was estimated using a BioSpec-nano (Shimadzu). The RNA samples were then stored at −80°C. RNA libraries and sequencing were performed using the Illumina NovaSeq6000 platform.

### Gene expression analysis

The sequence quality of the resulting raw Illumina reads was assessed with FastQC (v0.11.9) and processed in Trimmomatic 0.39 with the parameters ILLUMINACLIP: TruSeq3-PE.fa:2:30:10 LEADING:30 TRAILING:30 SLIDINGWINDOW:4:20 MINLEN:101. All reads for each sample were aligned to the *D. suzukii* reference genome (GCF_013340165.1 from NCBI RefSeq assembly) using HiSat2 (Kim et al. 2015) to estimate gene expression levels. The alignment files were converted to BAM and sorted using SAMtools (Li et al. 2009). Effective counts were computed using StringTie (Pertea et al. 2015), and raw read count values were obtained using the Python script (prepDE.py). The read count data were used for gene expression analysis.

Homologs of *D. suzukii* genes were searched using BLASTx searches for *D. melanogaster* protein sequences (GCF_000001215.4 from NCBI RefSeq assembly). Genes with the best hit and an *e*-value < 0.0001 were used for the analysis. Differentially expressed genes (DEGs) among light treatment, urbanization type, and the interaction between these factors were detected using DESeq2 (Love et al. 2014). Genes with *q* < 0.05 and fold change > 1.5 were considered as DEGs. Gene ontology (GO) enrichment analysis of the DEGs was performed, and the GO terms were simplified by similarity using the R package ‘clusterProfiler’ (Wu et al. 2021).

### Statistical analyses

All statistical analyses were performed in R (v3.5.0). The effect of ALAN treatment on body size, principal component 1(PC1) and PC2 of activity level, the length of the circadian rhythm period, and PC1 and PC2 of gene expression were analyzed. The effect of ALAN on the differences in these parameters between urban and rural populations was also analyzed using a linear mixed model (LMM) with the *lmer* function from the ‘lme4’ package. The model was constructed separately for wing length, thorax breadth, PC1 and PC2 of activity level, and length of the circadian rhythm period as response variables. The main effects of light treatment (L: control or ALAN), urbanization type (U: urban or rural), sex (S: male or female), and their two-way and three-way interaction effects were included as predictors. PC1 and PC2 of gene expression were also analyzed separately for males and females. The effect of ALAN treatment on the sleep fraction and its differences between urban and rural populations were analyzed using LMM, with the sleep fraction as the response variable. In the model, the main effects of L, U, S, time of day (T: day or night), and their two-way interaction effects were included. A random intercept for the isofemale line was included to account for nonindependence among flies from the same urbanization type.

## Results

The wings and thorax sizes were larger in females than in males, regardless of the urbanization type and light treatment (wing: χ^2^ = 177.7, *P* < 0.001; thorax: χ^2^ = 110.3, *P* < 0.001; Fig. 1A). Exposure to ALAN significantly decreased wing and thorax size (wing: *χ*^2^ = 11.81, *P =* 0.001; thorax: χ^2^ = 4.0, *P =* 0.046). The wing size of the urban individuals was significantly larger than that of the rural ones (*χ*^2^ = 5.09, *P =* 0.0241). No significant interaction effect was observed for wing size (L × U: χ^2^ = 0.53, *P* = 0.47; L × S: χ^2^ = 0.24, *P =* 0.62; S × U: χ^2^ = 0.01, *P =* 0.92; L × S × U: χ^2^ = 0.23, *P =* 0.63) and thorax size (U: χ^2^ = 0.001, *P* = 0.97; L × U: χ^2^ = 0.55, *P* = 0.45; L × S: χ^2^ = 0.012, *P =* 0.91; S × U: χ^2^ = 0.20, *P =* 0.65; L × S × U: χ^2^ = 0.11, *P =* 0.75). However, the reduction in wing and thorax size in females in response to ALAN exposure tended to be smaller in individuals from urban populations than those from rural populations.

**Figure 1.**
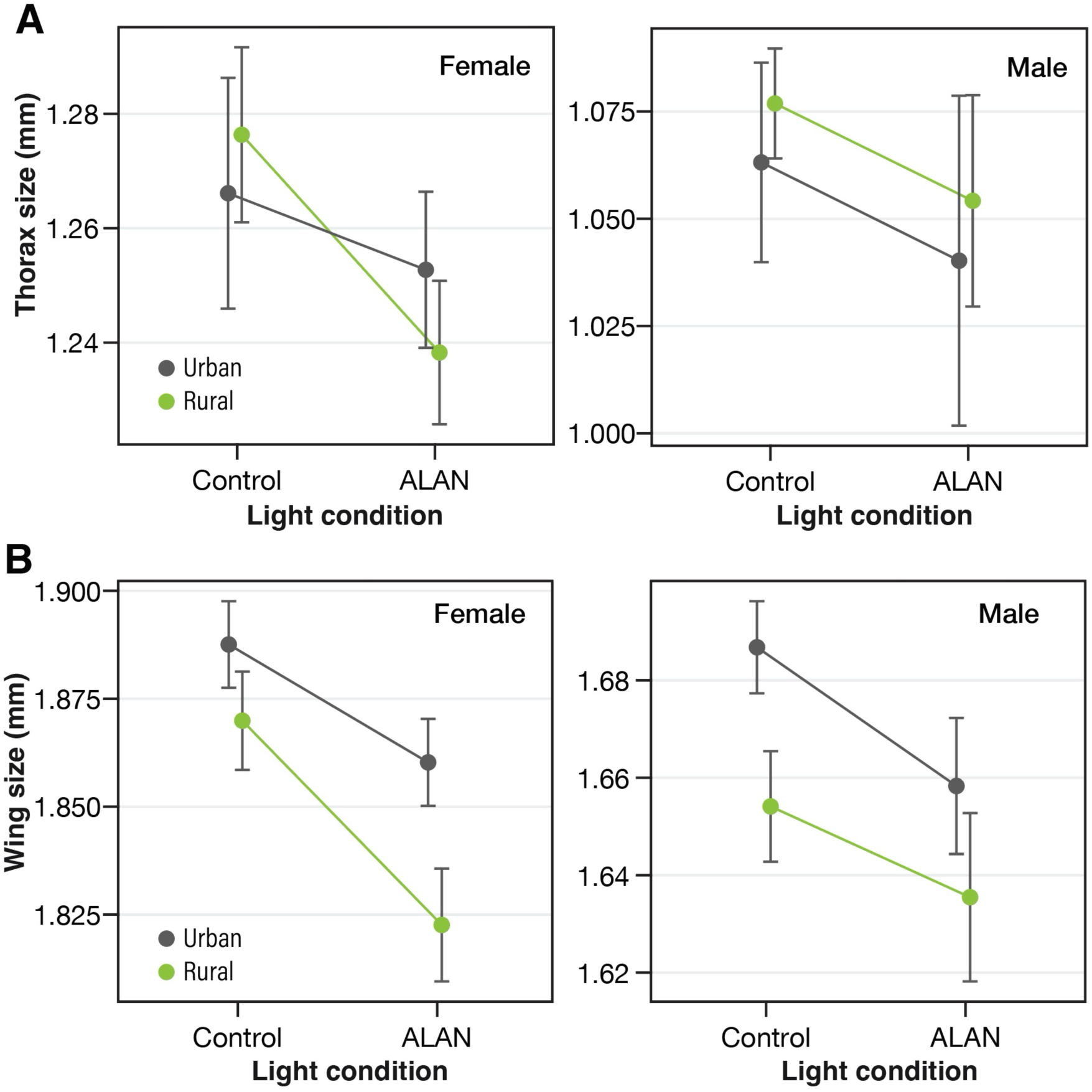
Effect of artificial light at night on the body size of individuals from urban and rural populations. (A) Thorax length and (B) wing length.

The maximum survival time was 85 days (Fig. 2). The median time for the control and ALAN-treated groups in the urban population was 28 and 58 days, respectively; it was 42 and 41 days, respectively, for the rural population. The main effects of sex, urbanization type, and light treatment did not explain the variation in the survival rate (S: *z* = 0.18, *P =* 0.86; U: *z* = 0.57, *P =* 0.57; L: *z* = 0.61, *P =* 0.54; see Table S2). A significant interaction effect between urbanization type and light treatment on survival rate was observed (*z* = −2.17; *P* = 0.03), indicating that rural populations tended to survive longer under the control condition than under ALAN treatment, while urban populations survived longer under ALAN treatment than under the control condition.

**Figure 2.**
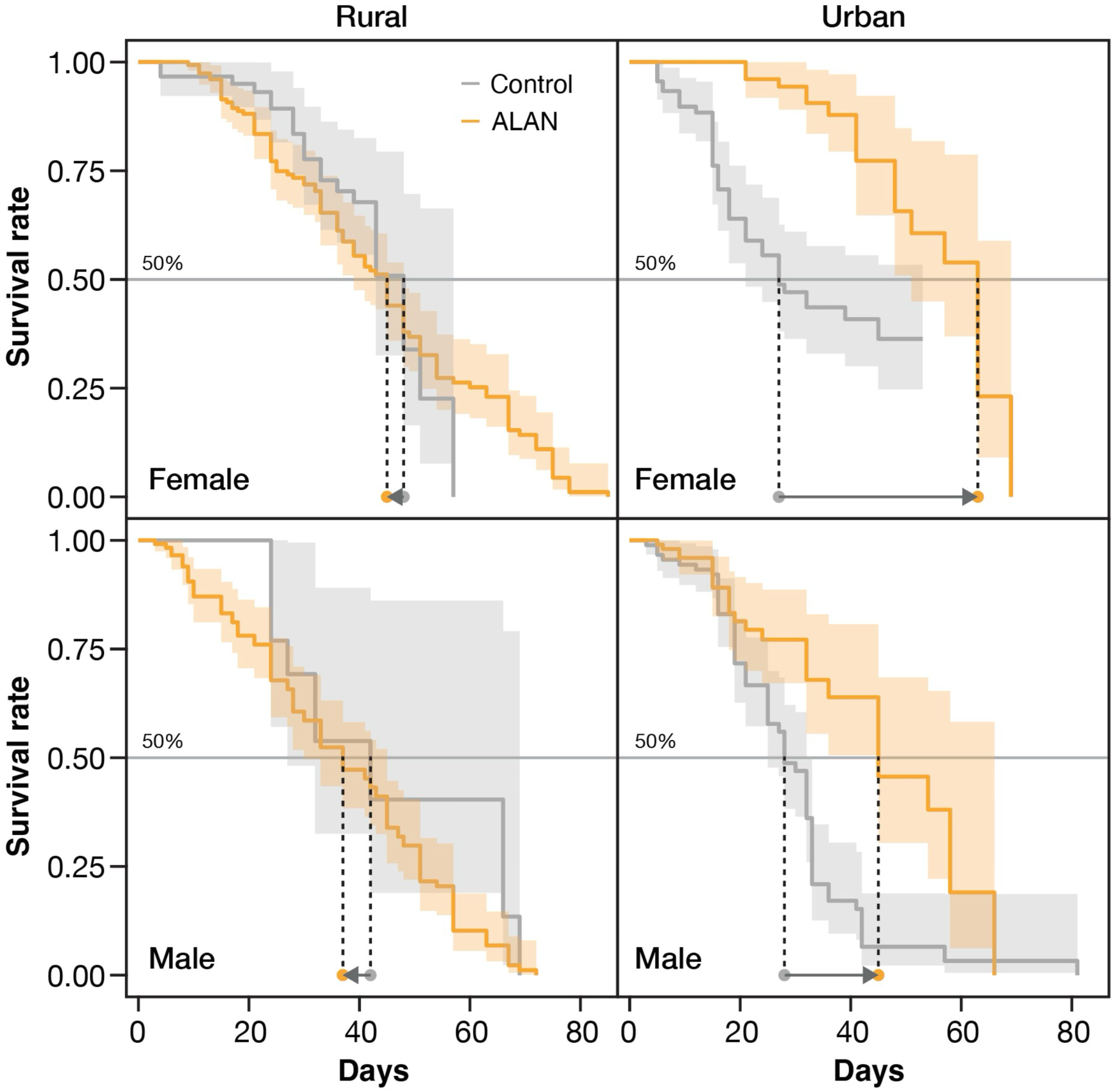
Survival curves of flies under control and ALAN conditions. The dotted lines represent the estimated median, which is the age at which the curves intersect at 50%.

Locomotor activities under the dark condition displayed a monophasic pattern, peaking at dusk (Fig. 3A). This peak tended to shift to late night under the ALAN treatment. The activity level during nighttime (shaded time zone in Fig. 3A) was quite low. The daily activity patterns on the second day were similar to those on the first day, although the activity level was relatively low. As a result of PCA with data for the first day, two central PCs were identified (Fig. S1). PC1 explained 23.6% of the variance and was derived from the total amount of activity (i.e., activity level). A lower PC1 value indicated a higher activity level (Fig. 3B). PC2, which explained 14.4% of the variance, reflected the relative strength of the activity level around zeitgeber time 10–18, corresponding to twilight time and early night, compared to daytime (Fig. 3B). A higher PC2 value indicated a higher activity level at twilight and early night. The variances of the other PCs were small (< 5%). The PC1 score of individuals exposed to ALAN was lower than that of the control group (L: χ^2^ = 56.4, *P* < 0.001), indicating a higher activity level under ALAN treatment. Males were significantly more active per day, but no differences were observed in activity by urbanization type (U: χ^2^ = 0.13, *P =* 0.72; S: χ^2^ = 230.7, *P* < 0.001). No significant interactions, including those between urbanization type and light treatment, were observed (L × U: χ^2^ = 0.20, *P* = 0.65; L × S: χ^2^ = 0.46, *P =* 0.50; S × U: χ^2^ = 0.08, *P =* 0.78; L × S × U: χ^2^ = 0.05, *P =* 0.82), indicating that the response to ALAN was the same in urban and rural populations. The PC2 scores of male individuals exposed to ALAN were higher than those of control individuals in rural and urban populations. In contrast, the PC2 scores of urban females exhibited little sensitivity to light conditions compared to others. An interaction effect between sex and light treatment and a three-way interaction for PC2 (L × S: χ^2^ = 6.05, *P* = 0.014; L × S × U: χ^2^ = 5.83, *P* = 0.016; Fig. 3B) were observed. No significant main effects of urbanization type on PC2 and no interaction effects between type and light treatment or type and sex were identified (U: χ^2^ = 0.39, *P* = 0.53; L × U: χ^2^ = 1.84, *P* = 0.18; S × U: χ^2^ = 1.40, *P =* 0.24).

**Figure 3.**
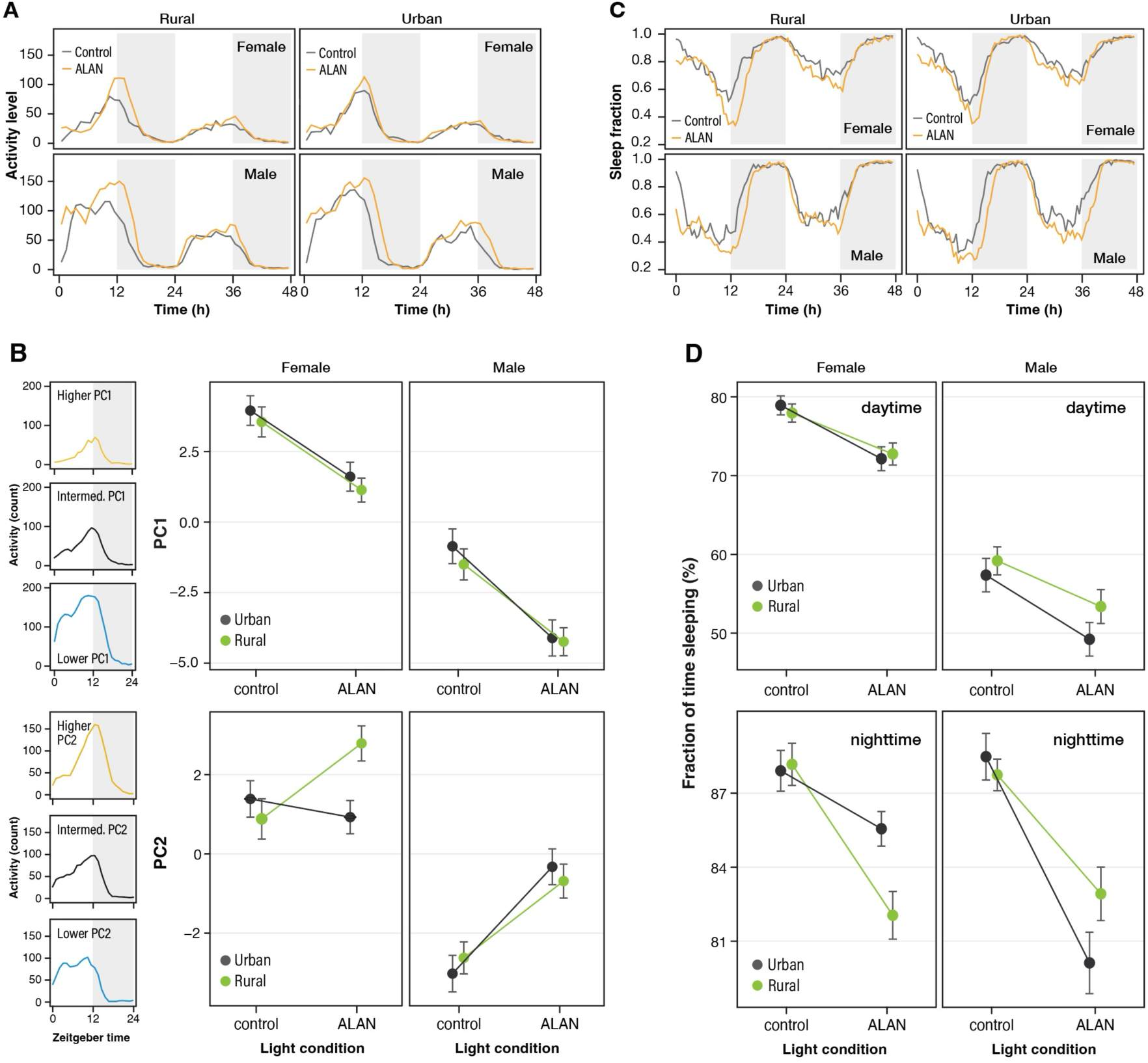
Effects of ALAN on activity patterns of urban and rural populations. (A) Each population’s locomotor activity counts per hour under control and ALAN treatments. (B) Bright gray shading indicates the potential daytime under DD conditions, and darker gray shading indicates the potential nighttime. Biplot in PC1 and PC2 space (left panel). Individuals’ daily physical activity patterns were divided into three categories according to the PC values: PC1 (center panel) and PC2 (right panel). (C) Relationship between light condition and PC1 or PC2. (D) Fraction of time spent sleeping averaged within a 30-min time window (D). (E) Relationship between the light condition and the fraction of time spent sleeping.

Figure 3C presents the daily pattern of sleep behavior. The sleep fraction was higher at nighttime than at daytime, regardless of treatment (χ^2^ = 0.20, *P* < 0.001). The sleep onset time was later under the ALAN treatment than the control treatment. Sleep time during daytime and nighttime was reduced under the ALAN treatment (χ^2^ = 131.2, *P* < 0.001). No significant difference by urbanization type was observed, but a main effect of sex and interactions between sex and time and sex and type was revealed (S: χ^2^ = 474.3, *P* < 0.001; T × S: χ^2^ = 254.3, *P* < 0.001; P × S: χ^2^ = 8.57, *P* = 0.0034). No significant main effect of urbanization type and no other interactions on sleep duration were found (U: χ^2^ = 0.02, *P* = 0.89; L × T: χ^2^ = 0.84, *P* = 0.66; L × U: χ^2^ = 1.18, *P* = 0.28; L × S: χ^2^ = 3.21, *P* = 0.07; T × U: χ^2^ = 1.93, *P* = 0.38).

The length of the endogenous rhythms was shortened by ALAN treatment (S: χ^2^ = 70.4, *P* < 0.001; U: χ^2^ = 0.0004, *P* = 0.98; L: χ^2^ = 7.37, *P* = 0.0067; Fig. 4). A reduced length of endogenous rhythms was observed in individuals from rural populations, and the interaction between urbanization type and light treatment was significant (L × U: χ^2^ = 5.25, *P* = 0.022; L × S: χ^2^ = 2.93, *P* = 0.087; P × S: χ^2^ = 0.04, *P* = 0.84; L × S × U: χ^2^ = 0.09, *P* = 0.76).

**Figure 4.**
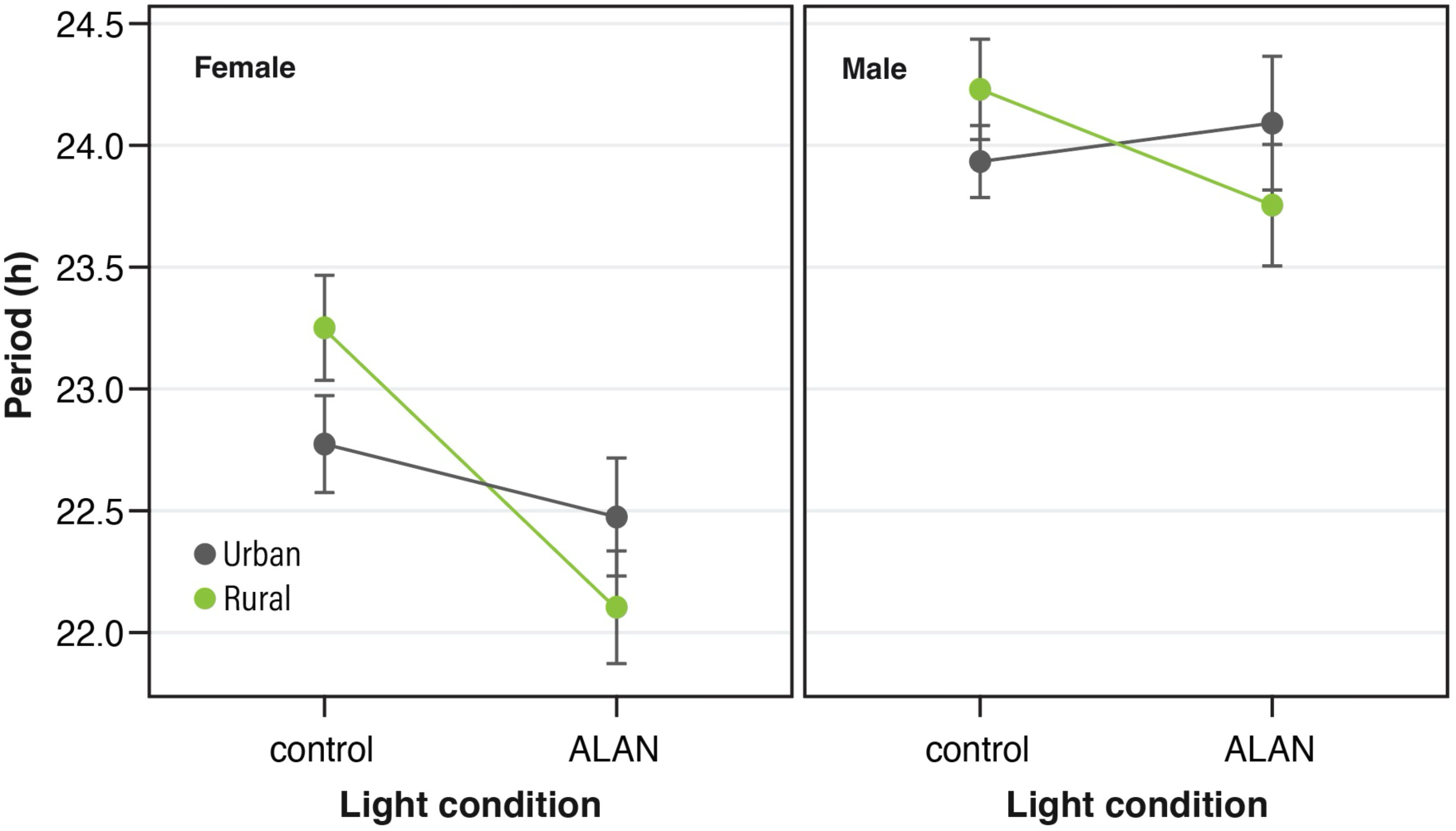
Effects of ALAN on the circadian rhythm period of individuals from urban and rural populations.

Among the 45,488 transcripts assembled, 37,895 (83.3%) were annotated to 20,505 genes. After removing transcripts whose read counts among all samples were zero, 18,754 transcripts were used for downstream analyses. Figure 5A presents the outcome of PCA using the gene expression data. The changes in the gene expression profile due to ALAN exposure were not clear in males. In contrast, the expression profile of females tended to be changed clearly by ALAN exposure, especially in urban individuals. For PC1, while the effect of the sex on PC scores was significant (χ^2^ = 873.0, *P* < 0.001), the effect of ALAN exposure and type (L: χ^2^ = 3.70, *P* = 0.054; U: χ^2^ = 2.93, *P* = 0.087), the two-way interaction effect among ALAN exposure, sex, and type (L × U: χ^2^ = 0.37, *P* = 0.54; L × S: χ^2^ = 2.62, *P* = 0.11; P × S: χ^2^ = 0.37, *P* = 0.54), and the three-way interaction effect of them on PC scores were not significant (L × S × U: χ^2^ = 1.26, *P* = 0.26). For PC2, while the effect of the sex (χ^2^ = 10.9, *P* < 0.001) and the three-way interaction effect of ALAN, sex, and urbanization type on PC scores were significant (χ^2^ = 6.36, *P* = 0.012), the main effect of ALAN exposure and type (L: χ^2^ = 0.85, *P* = 0.36; U: χ^2^ = 0.44, *P* = 0.51) and the two-way interaction of effect among ALAN, sex, and type on PC scores were not significant (L × U: χ^2^ = 0.15, *P* = 0.70; L × S: χ^2^ = 0.39, *P* = 0.53; P × S: χ^2^ = 0.06, *P* = 0.81; Fig. 5B). When males and females were separately subjected to statistical analysis, the effect of ALAN on PC1 was significant for females but not for males (Tables S3 and S4). Likewise, the two-way interaction effect of ALAN and urbanization type on PC2 was significant for females but not for males (Tables S5 and S6). Thus, females derived from the urban population displayed a larger shift in the expression profile than those from the rural population.

**Figure 5.**
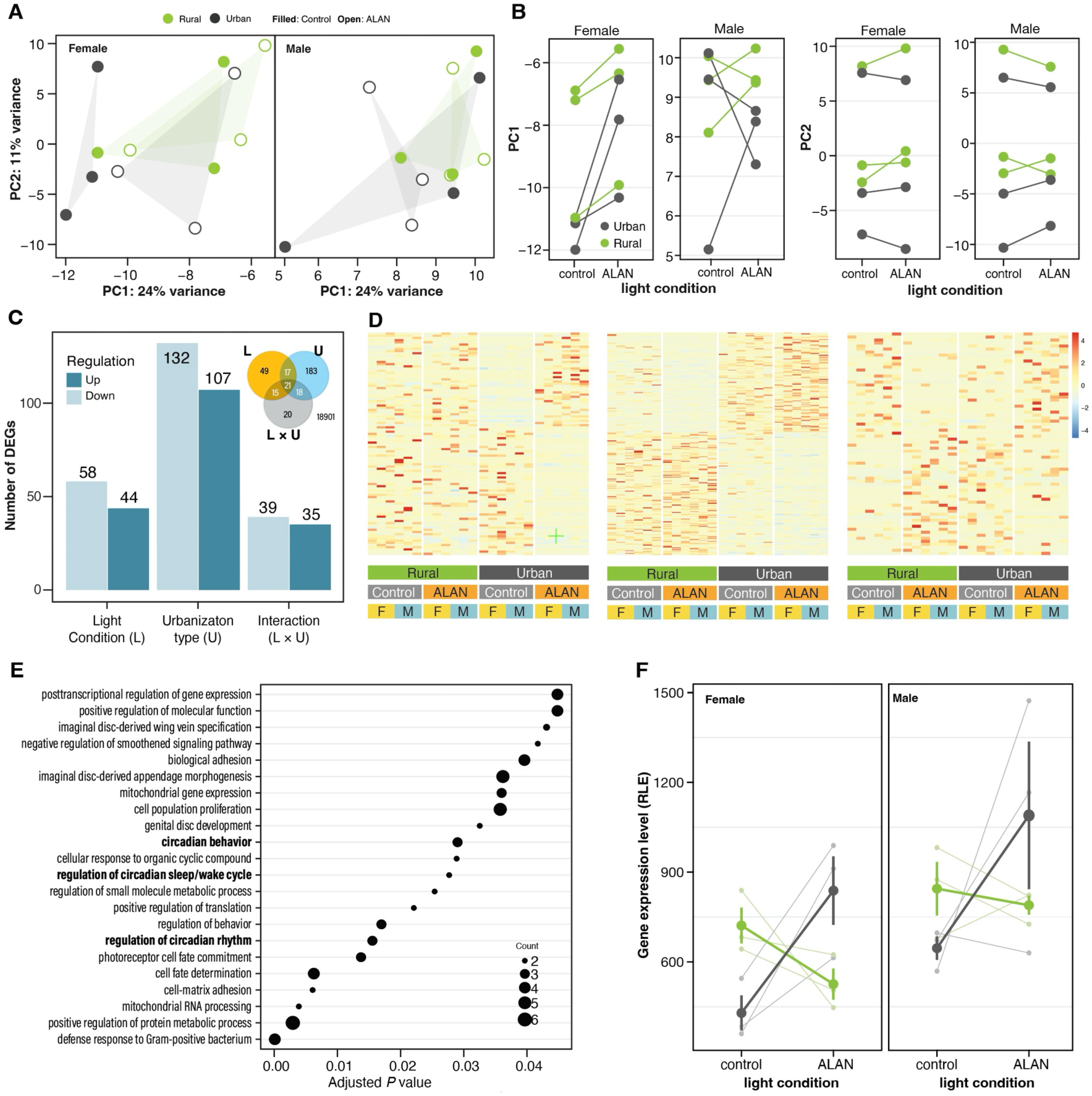
Effects of ALAN on the gene expression of individuals from urban and rural populations. (A) PCA plots representing changes in the gene expression profile with ALAN exposure. Each point represents an independent biological replicate. (B) Changes in PC1 and PC2 scores with ALAN exposure. (C) Number of DEGs. (D) Heatmap of DEGs for light treatments, urbanization types, and their interaction. (E) GO terms for BP enriched in DEGs of the interaction between light treatment and urbanization type. (F) Effects of ALAN on the gene expression level of *LARK* in individuals of urban and rural populations. Small plots and thin lines represent the expression level for each population.

The expression levels of 18,754 genes were analyzed, and these genes were classified into 3 categories based on their expression variation patterns:107 genes were upregulated, while 132 genes were downregulated in urban individuals compared to rural populations (Fig. 5C, D). Forty-four genes were significantly upregulated, while 58 genes were significantly downregulated under ALAN exposure. Seventy-four genes were significantly influenced by the interaction between urbanization type and light treatments, suggesting that their expression levels responded differently to ALAN in urban and rural populations. DEGs of these three categories partly overlapped (inset figure of Fig. 5C). In the GO enrichment analysis for the DEGs of the three categories, no GO terms for biological process (BP) were detected for the DGEs induced by ALAN, while the GO term “photoreceptor cell fate commitment” was detected for the DEGs relating to the urbanization type. On the other hand, 22 GO terms for BP were obtained in the DEGs that were affected by the interaction effect between urbanization type and light treatment (Fig. 5E). The GO terms included circadian rhythm-related categories, such as “circadian behavior,” “regulation of circadian sleep/wake cycle,” and “regulation of circadian rhythm,” in addition to “photoreceptor cell fate commitment.” For instance, the expression of *LARK*, identified as a circadian rhythm-related DEG influenced by the interaction of urbanization type and light treatment, decreased under ALAN in rural populations but increased in urban populations (L: P = 0.07, S: *P* < 0.05, L × U: *P* < 0.001).

## Discussion

ALAN has spread rapidly due to urbanization, contributing to environmental changes in urban areas. Recent studies have revealed that various species persist in cities through phenotypic plasticity or adaptive evolution in response to urbanization stress. However, most studies have failed to distinguish between nongenetic and genetic changes. This study examined the effects of ALAN on body size, survival, activity patterns, gene expression, and evolutionary responses to ALAN in the urban populations of *D. suzukii*. Laboratory experiments revealed that ALAN alters a range of traits and behaviors. Furthermore, our results reveal that urban populations have undergone adaptive evolution to withstand ALAN. Evolutionary adaptations to urban stressors may have contributed to the successful colonization of urban areas by these species. *D. suzukii* exhibits a bimodal activity pattern peaking at dawn and dusk, with concentrated sleep at night (Evans et al. 2017). The unimodal activity observed in this study may be induced by the lack of zeitgeber signals. Indeed, the morning activity peak tends to disappear under DD conditions in *D. melanogaster* (Dubruille and Emery 2008). Additionally, ALAN increased the overall activity and shifted the peak of activity at dusk, leading to an increase in activity levels in the early night. A similar change in the activity pattern due to ALAN exposure was reported in a previous study under LD conditions (Sato and Takahashi 2022). ALAN may induce unusual activity during the night. Consequently, ALAN also decreased the sleep fraction. It tended to delay the peak of activity and sleep in the evening. This pattern is consistent with a previous study reporting that ALAN shifts dusk activity peaks to nighttime and increases average activity levels in *D. melanogaster* (Kempinger et al. 2009). ALAN may disrupt the diurnal activity of this species, potentially having adverse effects on its development and survival. Additionally, our study revealed that ALAN shortens the circadian clock. Generally, a mismatch between endogenous and environmental rhythm lengths harms growth and survival (Ouyang et al. 1998). Adaptations to match endogenous cycles with environmental rhythms increase fitness in diverse taxa (Dodd et al. 2005; Horn et al. 2019). These facts suggest that the ALAN-induced shortening of endogenous rhythms also influences development and survival, harming fitness. Importantly, the females of urban populations did not display an unusual increase in nighttime activity, and the effect of ALAN on the circadian rhythm length was not observed in the urban population. These findings indicate that urban populations have developed resilience to ALAN stress, revealing the presence of contemporary evolution in urban environments.

Generally, a positive correlation between body (wing) size and dispersal ability is observed; larger body size is considered adaptive in urban areas where habitats are highly fragmented (Piano et al. 2017). The larger wing size of urban individuals compared to rural ones is consistent with previous studies on urban insect populations (San Martin et al. 2012; Schoville et al. 2013). Additionally, longer flight seasons in warmer regions, including urban areas, promote larger body sizes and slower development (Masaki 1967; Mousseau 1997). Larger wings in urban populations under control conditions indicate adaptive evolution to urban areas with fragmented habitats and extended flight seasons. In this context, decreased wing and thorax sizes caused by ALAN exposure are unlikely to represent an adaptive response to the urban environment. On the other hand, previous studies have demonstrated that ALAN can shorten or extend the insects’ developmental period, resulting in a reduced or increased body size (Van Geffen et al. 2014; Durrant et al. 2018b). The mechanism linking ALAN to body size reduction remains unclear; however, a decreased developmental period may contribute to the observed reduced body size.

The fitness impacts of size reduction caused by ALAN in urban environments remain a key topic for future research. Notably, our previous study reported that ALAN shifts *D. suzukii* toward an r-strategy, characterized by producing many small offspring (Sato and Takahashi 2024). In any case, our findings indicate a reduced body size response to ALAN stress in urban populations, particularly females, suggesting adaptive evolution for ALAN tolerance. This result is further supported by observed changes in daily activity patterns.

The shorter life spans of rural individuals exposed to ALAN align with a previous study revealing that *D. melanogaster* adults exposed to 10 Lx and 100 Lx died earlier than those exposed to 0 Lx (McLay et al. 2017). In contrast, ALAN-treated urban individuals lived longer than those under control conditions, suggesting that urban populations may have evolved to tolerate or even benefit from ALAN exposure. The present study suggests that ALAN could affect development and survival and that urban *D. suzukii* populations may overcome ALAN stress in urban environments. However, at this time, it was not possible to determine whether the effects of ALAN experienced during the larval stage or the adult stage influenced adult survival time. Since ALAN exposure during the larval stage alters body size and reproductive investment in *D. suzukii* (Sato and Takahashi 2024), the effects of ALAN during the larval stage alone could influence adult survival time. It would be necessary to measure survival time by switching rearing conditions between the larval and adult stages to conduct a more detailed investigation of the effects of ALAN.

The transcriptome analysis revealed that ALAN had significant effects on the gene expression profile. Changes in the expression levels of certain genes differed between individuals from rural and urban populations. These genes were functionally associated with photoreceptors and circadian rhythms, as previously reported in studies on ALAN’s effects on gene expression in firefly larvae (Chen et al. 2021). One such gene, *LARK*, functions as a translational regulator involved in circadian cycle regulation, and changes in *LARK* abundance can alter the circadian cycle (Price 2014). Rural populations with decreased *LARK* expression exhibited shorter circadian cycle lengths than urban populations with significantly increased *LARK* expression, which displayed smaller ALAN-induced changes in the circadian cycle length. Evolutionary changes in the responsiveness of circadian rhythm-related genes to ALAN may contribute to the robustness of diurnal rhythms in urban populations, reducing the phenotypic negative impact of nighttime light exposure.

Furthermore, sex differences in response to ALAN exposure were observed as significant two-way and three-way interaction effects involving sex and light treatment for various traits. Typically, the influence of ALAN on phenotypic traits appeared more ambiguous in urban females than in rural females or males from either population. Interestingly, changes in gene expression patterns were more pronounced in urban females than in rural females and males from either population. The contrasting patterns between phenotypes and gene expression observed in this study suggest that the pronounced changes in gene expression in urban females in response to ALAN could help mitigate its effects at the phenotypic level, improving tolerance to urban stress. The evolution of adaptive plasticity in gene expression may contribute to the adaptation of these species to urban environments (Campbell-Staton et al. 2021).

We examined the effects of ALAN on the phenotypes, gene expression, and evolutionary divergence between the *D. suzukii* urban and rural populations. Phenotypic analyses indicate that ALAN affects growth, development, and activity, with urban populations exhibiting greater resilience to urban stress than their rural counterparts. However, gene expression patterns, especially in females, were more sensitive in urban individuals. These findings suggest that the resilience of urban individuals may stem from enhanced plasticity in their gene expression patterns. Further studies are required to examine the influence of ALAN on various traits, including reproduction, and compare nucleotide sequences in regulatory regions between urban and rural populations to fully elucidate the effects of ALAN on organisms and the molecular mechanisms underlying evolutionary responses.

## Acknowledgements

The research was funded the Asahi Group Foundation, The Japan Prize Foundation, Obayashi Foundation and The Sumitomo Foundation. This research was performed by the Environment Research and Technology Development Fund (4RF-2103) of the Environmental Restoration and Conservation Agency of Japan.

## Author contributions

N.T. and Y.T. developed the general design of the study, the experiments, and analyses. N.T. performed the experiment, carried out the statistical analyses with input from Y.T. and wrote the first version of the manuscript. Y.T. edited the first version, and N.T. and Y.T. contributed equally to the further editing of subsequent versions of the manuscript.

## Data availability

Data are available online (figshare: *******).

## Supplemental Information

**Table S1.** Populations used in the experiments.

| Population | Latitude | Longitude | Urbanization type | Experiment |
| --- | --- | --- | --- | --- |
| ITA | 35.3206000 | 140.1423000 | rural | BS, LA, GE |
| RES | 35.4784570 | 140.2439530 | rural | BS, LA, GE |
| INO | 35.7002000 | 139.5764000 | urban | BS, SR, LA, GE |
| SUM | 35.7126000 | 139.8040000 | urban | BS, SR, LA |
| YAY | 35.6287000 | 140.1026000 | urban | SR, LA |
| NOG | 35.4631000 | 140.1129000 | rural | SR, LA, GE |
| KSI | 35.6407740 | 139.8553310 | urban | SR, LA |
| KUR | 35.2877140 | 140.0898190 | rural | SR, LA |
| TEN | 35.7056890 | 140.5477960 | rural | SR, LA |
| TKB | 36.2206880 | 140.1051640 | rural | SR, LA |
| WAK | 35.6755870 | 139.9936830 | urban | SR, LA |
| ASU | 35.7504718 | 139.7366288 | urban | SR, LA, GE |
| TTY | 34.9803920 | 139.8555450 | rural | SR, LA |
| STB | 35.6936817 | 139.7385036 | urban | SR, LA, GE |
| IZU | 35.5789340 | 140.2268830 | rural | LA |
| SEA | 35.6093087 | 139.7474079 | urban | LA |

BS, test for body size; SR, test for survival rate; LA, test for locomotor activity; GE, test for gene expression.

**Table S2.** Results of the Cox mixed-effects model for the survival rate.

| Factors | *Z* | *P* |
| --- | --- | --- |
| light treatment | 0.61 | 0.54 |
| urbanization type | 0.57 | 0.57 |
| sex | 0.18 | 0.86 |
| light treatment × urbanization type | −2.17 | 0.03 |
| light treatment × sex | 0.36 | 0.72 |
| urbanization type × sex | 0.01 | 0.99 |
| light treatment × urbanization type × sex | 0.06 | 0.95 |

**Table S3.** Results of LMM for the PC1 valued of principal component analysis of gene.

| Factors | $\chi^2$ | *P* |
| --- | --- | --- |
| Light treatment | 0.044 | 0.83 |
| Urbanization type | 2.1 | 0.15 |
| Light treatment × urbanization type | 0.13 | 0.72 |

**Table S4.** Results of LMM for the PC1 values of PCA of gene expression in females.

| Factors | $\chi^2$ | *P* |
| --- | --- | --- |
| Light treatment | 13.0 | 0.00030 |
| Urbanization type | 1.9 | 0.17 |
| Light treatment × urbanization type | 3.1 | 0.078 |

**Table S5.** Results of LMM for the PC2 values of PCA of gene expression in males.

| Factors | $\chi^2$ | *P* |
| --- | --- | --- |
| Light treatment | 0.022 | 0.88 |
| Urbanization type | 0.45 | 0.50 |
| Light treatment × urbanization type | 1.2 | 0.28 |

**Table S6.** Results of LMM for the PC2 values of PCA of gene expression in females.

| Factors | $\chi^2$ | *P* |
| --- | --- | --- |
| Light treatment | 1.4 | 0.24 |
| Urbanization type | 0.44 | 0.51 |
| Light treatment × urbanization type | 4.9 | 0.026 |

## References

Campbell-Staton, S. C., J. P. Velotta, and K. M. Winchell. 2021. Selection on adaptive and maladaptive gene expression plasticity during thermal adaptation to urban heat islands. Nat Commun 12. Nature Research.

Chen, Y. R., W. L. Wei, D. T. W. Tzeng, A. C. S. Owens, H. C. Tang, C. S. Wu, S. S. Lin, S. Zhong, and E. C. Yang. 2021. Effects of artificial light at night (ALAN) on gene expression of Aquatica ficta firefly larvae. Environmental Pollution 281. Elsevier Ltd.

Cinzano, P., F. Falchi, and C. D. Elvidge. 2001. The first World Atlas of the artificial night sky brightness. Mon Not R Astron Soc 328:689–707.

Dodd, A. N., N. Salathia, A. Hall, E. Kévei, R. Tóth, F. Nagy, J. M. Hibberd, A. J. Millar, and A. A. R. Webb. 2005. Plant Circadian Clocks Increase Photosynthesis, Growth, Survival, and Competitive Advantage. Science (1979) 309:630–633.

Dubruille, R., and P. Emery. 2008. A plastic clock: How circadian rhythms respond to environmental cues in drosophila. Humana Press.

Durrant, J., L. M. Botha, M. P. Green, and T. M. Jones. 2018a. Artificial light at night prolongs juvenile development time in the black field cricket, Teleogryllus commodus. J Exp Zool B Mol Dev Evol 330:225–233. John Wiley and Sons Inc.

Durrant, J., L. M. Botha, M. P. Green, and T. M. Jones. 2018b. Artificial light at night prolongs juvenile development time in the black field cricket, *Teleogryllus commodus*. J Exp Zool B Mol Dev Evol 330:225–233.

Durrant, J., E. B. Michaelides, T. Rupasinghe, D. Tull, M. P. Green, and T. M. Jones. 2015. Constant illumination reduces circulating melatonin and impairs immune function in the cricket Teleogryllus commodus. PeerJ 2015. PeerJ Inc.

Evans, R. K., M. D. Toews, and A. A. Sial. 2017. Diel periodicity of Drosophila suzukii (Diptera: Drosophilidae) under field conditions. PLoS One 12. Public Library of Science.

Falchi, F., P. Cinzano, D. Duriscoe, C. C. M. Kyba, C. D. Elvidge, K. Baugh, B. A. Portnov, N. A. Rybnikova, and R. Furgoni. 2016. The new world atlas of artificial night sky brightness. Sci Adv 2. American Association for the Advancement of Science.

Fukano, Y., W. Yamori, H. Misu, M. P. Sato, K. Shirasawa, Y. Tachiki, and K. Uchida. 2023. From green to red: Urban heat stress drives leaf color evolution. Sci Adv 9.

Geissmann, Q., L. G. Rodriguez, E. J. Beckwith, and G. F. Gilestro. 2019. Rethomics: An R framework to analyse high-throughput behavioural data. PLoS One 14. Public Library of Science.

Hahs, A. K., B. Fournier, M. F. J. Aronson, C. H. Nilon, A. Herrera-Montes, A. B. Salisbury, C. G. Threlfall, C. C. Rega-Brodsky, C. A. Lepczyk, F. A. La Sorte, I. MacGregor-Fors, J. Scott MacIvor, K. Jung, M. R. Piana, N. S. G. Williams, S. Knapp, A. Vergnes, A. A. Acevedo, A. M. Gainsbury, A. Rainho, A. J. Hamer, A. Shwartz, C. C. Voigt, D. Lewanzik, D. M. Lowenstein, D. O’Brien, D. Tommasi, E. Pineda, E. S. Carpenter, E. Belskaya, G. L. Lövei, J. C. Makinson, J. L. Coleman, J. P. Sadler, J. Shroyer, J. T. Shapiro, K. C. R. Baldock, K. Ksiazek-Mikenas, K. C. Matteson, K. Barrett, L. Siles, L. F. Aguirre, L. O. Armesto, M. Zalewski, M. I. Herrera-Montes, M. K. Obrist, R. K. Tonietto, S. A. Gagné, S. J. Hinners, T. Latty, T. D. Surasinghe, T. Sattler, T. Magura, W. Ulrich, Z. Elek, J. Castañeda-Oviedo, R. Torrado, D. J. Kotze, and M. Moretti. 2023. Urbanisation generates multiple trait syndromes for terrestrial animal taxa worldwide. Nat Commun 14. Nature Research.

Hoffmann, J., R. Palme, and J. A. Eccard. 2018. Long-term dim light during nighttime changes activity patterns and space use in experimental small mammal populations. Environmental Pollution 238:844–851. Elsevier Ltd.

Horn, M., O. Mitesser, T. Hovestadt, T. Yoshii, D. Rieger, and C. Helfrich-Förster. 2019. The circadian clock improves fitness in the fruit fly, Drosophila melanogaster. Front Physiol 10. Frontiers Media S.A.

Huang, K., X. Li, X. Liu, and K. C. Seto. 2019. Projecting global urban land expansion and heat island intensification through 2050. Environmental Research Letters 14. Institute of Physics Publishing.

Jiang, J., Y. He, H. Kou, Z. Ju, X. Gao, and H. Zhao. 2020. The effects of artificial light at night on Eurasian tree sparrow (Passer montanus): Behavioral rhythm disruption, melatonin suppression and intestinal microbiota alterations. Ecol Indic 108. Elsevier B.V.

Johnson, M. T. J., and J. Munshi-South. 2017. Evolution of life in urban environments. Science (1979) 358:eaam8327.

Karen Ho, B. S., and A. Sehgal. 2000. *Drosophila melanogaster*: An Insect model for fundamental studies of sleep.

Kempinger, L., R. Dittmann, D. Rieger, and C. Helfrich-Förster. 2009. The nocturnal activity of fruit flies exposed to artificial moonlight is partly caused by direct light effects on the activity level that bypass the endogenous clock. Chronobiol Int 26:151–166.

Kim, D., B. Langmead, and S. L. Salzberg. 2015. HISAT: a fast spliced aligner with low memory requirements. Nat Methods 12:357–360.

Kolbe, J. J., H. A. Moniz, O. Lapiedra, and C. J. Thawley. 2021. Bright lights, big city: an experimental assessment of short-term behavioral and performance effects of artificial light at night on Anolis lizards. Urban Ecosyst 24:1035–1045.

Kyba, C. C. M., K. P. Tong, J. Bennie, I. Birriel, J. J. Birriel, A. Cool, A. Danielsen, T. W. Davies, P. N. den Outer, W. Edwards, R. Ehlert, F. Falchi, J. Fischer, A. Giacomelli, F. Giubbilini, M. Haaima, C. Hesse, G. Heygster, F. Hölker, R. Inger, L. J. Jensen, H. U. Kuechly, J. Kuehn, P. Langill, D. E. Lolkema, M. Nagy, M. Nievas, N. Ochi, E. Popow, T. Posch, J. Puschnig, T. Ruhtz, W. Schmidt, R. Schwarz, A. Schwope, H. Spoelstra, A. Tekatch, M. Trueblood, C. E. Walker, M. Weber, D. L. Welch, J. Zamorano, and K. J. Gaston. 2015. Worldwide variations in artificial skyglow. Sci Rep 5:8409.

Lack, J. B., M. J. Monette, E. J. Johanning, Q. D. Sprengelmeyer, and J. E. Pool. 2016. Decanalization of wing development accompanied the evolution of large wings in high-altitude Drosophila. Proc Natl Acad Sci U S A 113:1014–1019. National Academy of Sciences.

Lee, J. C., D. J. Bruck, A. J. Dreves, C. Ioriatti, H. Vogt, and P. Baufeld. 2011. In Focus: Spotted wing drosophila, *Drosophila suzukii*, across perspectives.

Li, H., B. Handsaker, A. Wysoker, T. Fennell, J. Ruan, N. Homer, G. Marth, G. Abecasis, and R. Durbin. 2009. The sequence alignment/map format and SAMtools. Bioinformatics 25:2078–2079.

Liu, Z., C. He, and J. Wu. 2016. The relationship between habitat loss and fragmentation during urbanization: An empirical evaluation from 16 world cities. PLoS One 11. Public Library of Science.

Love, M. I., W. Huber, and S. Anders. 2014. Moderated estimation of fold change and dispersion for RNA-seq data with DESeq2. Genome Biol 15. BioMed Central Ltd.

Masaki, S. 1967. Geographic variation and climatic adaptation in a field cricket (Orthoptera: Gryllidae). Evolution (N Y) 21:725–741.

McLay, L. K., M. P. Green, and T. M. Jones. 2017. Chronic exposure to dim artificial light at night decreases fecundity and adult survival in Drosophila melanogaster. J Insect Physiol 100:15–20.

Minias, P. 2023. The effects of urban life on animal immunity: Adaptations and constraints. Elsevier B.V.

Mousseau, T. A. 1997. Ectotherms follow the converse to Bergmann’s rule. Evolution (N Y) 51:630–632.

Multini, L. C., A. B. B. Wilke, and M. T. Marrelli. 2019. Urbanization as a driver for temporal wing-shape variation in Anopheles cruzii (Diptera: Culicidae). Acta Trop 190:30–36. Elsevier B.V.

Ouyang, Y., C. R. Andersson † ‡, T. Kondo, S. S. Golden, and C. Hirschie Johnson. 1998. Resonating circadian clocks enhance fitness in cyanobacteria.

Pertea, M., G. M. Pertea, C. M. Antonescu, T. C. Chang, J. T. Mendell, and S. L. Salzberg. 2015. StringTie enables improved reconstruction of a transcriptome from RNA-seq reads. Nat Biotechnol 33:290–295. Nature Publishing Group.

Piano, E., K. De Wolf, F. Bona, D. Bonte, D. E. Bowler, M. Isaia, L. Lens, T. Merckx, D. Mertens, M. van Kerckvoorde, L. De Meester, and F. Hendrickx. 2017. Urbanization drives community shifts towards thermophilic and dispersive species at local and landscape scales. Glob Chang Biol 23:2554–2564. Blackwell Publishing Ltd.

Price, J. L. 2014. Translational regulation of the Drosophila post-translational circadian mechanism. PLoS Genet 10. Public Library of Science.

Rivkin, L. R., J. S. Santangelo, M. Alberti, M. F. J. Aronson, C. W. de Keyzer, S. E. Diamond, M. J. Fortin, L. J. Frazee, A. J. Gorton, A. P. Hendry, Y. Liu, J. B. Losos, J. S. MacIvor, R. A. Martin, M. J. McDonnell, L. S. Miles, J. Munshi-South, R. W. Ness, A. E. M. Newman, M. R. Stothart, P. Theodorou, K. A. Thompson, B. C. Verrelli, A. Whitehead, K. M. Winchell, and M. T. J. Johnson. 2019. A roadmap for urban evolutionary ecology. Evol Appl 12:384–398. Wiley-Blackwell.

Russart, K. L. G., and R. J. Nelson. 2018. Light at night as an environmental endocrine disruptor. Elsevier Inc.

Salmón, P., A. Jacobs, D. Ahrén, C. Biard, N. J. Dingemanse, D. M. Dominoni, B. Helm, M. Lundberg, J. C. Senar, P. Sprau, M. E. Visser, and C. Isaksson. 2021. Continent-wide genomic signatures of adaptation to urbanisation in a songbird across Europe. Nat Commun 12. Nature Research.

San Martin y Gomez, G., and H. van Dyck. 2012. Ecotypic differentiation between urban and rural populations of the grasshopper Chorthippus brunneus relative to climate and habitat fragmentation. Oecologia 169:125–133.

Sato, A., and Y. Takahashi. 2022. Responses in thermal tolerance and daily activity rhythm to urban stress in Drosophila suzukii. Ecol Evol 12. John Wiley and Sons Ltd.

Sato, A., and Y. Takahashi. 2024. Urban–rural diversification in response to nighttime dim light stress in *Drosophila suzukii*. Biological Journal of the Linnean Society 143.

Schmidt, C., M. Domaratzki, R. P. Kinnunen, J. Bowman, and C. J. Garroway. 2020. Continent-wide effects of urbanization on bird and mammal genetic diversity. Proceedings of the Royal Society B: Biological Sciences 287. Royal Society Publishing.

Schoville, S. D., I. Widmer, M. Deschamps-Cottin, and S. Manel. 2013. Morphological clines and weak drift along an urbanization gradient in the butterfly, Pieris rapae. PLoS One 8. Public Library of Science.

Senar, J. C., M. J. Conroy, J. Quesada, and F. Mateos-Gonzalez. 2014. Selection based on the size of the black tie of the great tit may be reversed in urban habitats. Ecol Evol 4:2625–2632. John Wiley and Sons Ltd.

Theodorou, P., L. M. Baltz, R. J. Paxton, and A. Soro. 2021. Urbanization is associated with shifts in bumblebee body size, with cascading effects on pollination. Evol Appl 14:53–68.

Tüzün, N., L. Op de Beeck, and R. Stoks. 2017. Sexual selection reinforces a higher flight endurance in urban damselflies. Evol Appl 10:694–703. Wiley-Blackwell.

Van Geffen, K. G., R. H. A. Van Grunsven, J. Van Ruijven, F. Berendse, and E. M. Veenendaal. 2014. Artificial light at night causes diapause inhibition and sex-specific life history changes in a moth. Ecol Evol 4:2082–2089. John Wiley and Sons Ltd.

Vaz, S., S. Manes, G. Khattar, M. Mendes, L. Silveira, E. Mendes, E. de Morais Rodrigues, D. Gama-Maia, M. L. Lorini, M. Macedo, and P. C. Paiva. 2023. Global meta-analysis of urbanization stressors on insect abundance, richness, and traits. Elsevier B.V.

Wu, T., E. Hu, S. Xu, M. Chen, P. Guo, Z. Dai, T. Feng, L. Zhou, W. Tang, L. Zhan, X. Fu, S. Liu, X. Bo, and G. Yu. 2021. clusterProfiler 4.0: A universal enrichment tool for interpreting omics data. Innovation 2. Cell Press.

Xu, X., Y. Xie, K. Qi, Z. Luo, and X. Wang. 2018. Detecting the response of bird communities and biodiversity to habitat loss and fragmentation due to urbanization. Science of the Total Environment 624:1561–1576. Elsevier B.V.

